# Beta- and gamma-band oscillatory connectivity support naturalistic reading of continuous text

**DOI:** 10.1101/2023.08.21.554068

**Authors:** Jan Kujala, Sasu Mäkelä, Pauliina Ojala, Jukka Hyönä, Riitta Salmelin

## Abstract

Large-scale integration of information across cortical structures, building on neural connectivity, has been proposed to be a key element in supporting human cognitive processing. In electrophysiological neuroimaging studies of reading, quantification of neural interactions has been limited to the level of isolated words or sentences due to artefacts induced by eye movements. Here, we combined magnetoencephalography recording with advanced artefact rejection tools to investigate both cortico-cortical coherence and directed neural interactions during naturalistic reading of full-page texts. Our results show that reading vs. visual scanning of text was associated with wide-spread increases of cortico-cortical coherence in the beta- and gamma-bands. We further show that the reading task was linked with increased directed neural interactions compared to the scanning task across a sparse set of connections within a wide range of frequencies. Together, the results demonstrate that neural connectivity flexibly builds on different frequency bands to support continuous natural reading.

## 1 Introduction

Human cognitive processing including language perception and production has been thought to rely on more than just hierarchical progression of information and instead to build on multifaceted interactive neural systems (Mesulam, 1990). Yet, in time-resolved neuroimaging, most studies on language processing have focused on revealing the modulation of sequential neural activation patterns in distinct brain regions (Salmelin *et al*., 2000; Marinkovic *et al*., 2003; Hulten *et al*., 2019). However, during the last decades there has been increased interest in explicitly determining the role of neural interactions and large-scale integration of information (Singer, 1999; Palva *et al*., 2005; Salmelin & Kujala, 2006) which, mechanistically, has been proposed to be based on dynamic linkage across brain regions via phase-synchronization within multiple frequency bands (Singer & Gray, 1995; Varela *et al*., 2001; Fries, 2005). Using magnetoencephalography (MEG), in particular, numerous electrophysiological investigations have sought to address the architecture of neural connectivity regarding different cognitive processes such as attention (Gross *et al*., 2004; Doesburg *et al*., 2016; Lobier *et al*., 2018) and working memory (Palva *et al*., 2005; Palva, Monto, & Palva, 2010; Palva, Monto, Kulashekhar, *et al*., 2010; Watrous *et al*., 2013). Within the language domain, task-relevant modulation of inter-areal phase coupling has been observed for speech production (Korzeniewska *et al*., 2011; Liljeström, Kujala, *et al*., 2015; Liljeström, Stevenson, *et al*., 2015) and perception (Fonteneau *et al*., 2014; Saarinen *et al*., 2015), reading (Kujala *et al*., 2007, 2012; Perrone-Bertolotti *et al*., 2012; Vidal *et al*., 2012; Molinaro *et al*., 2013; Schoffelen *et al*., 2017; Liljeström *et al*., 2018) and writing (Saarinen *et al*., 2020). Such studies have typically focused on processing and perception of individual words (Korzeniewska *et al*., 2011; Kujala *et al*., 2012; Liljeström *et al*., 2018). Continuous, naturalistic stimulation has been used to examine connectivity during speech perception (Saarinen *et al*., 2015) but, in the case of reading, MEG studies of connectivity have been limited to sequential presentation of words, thus precluding eye movements associated with natural reading (Kujala *et al*., 2007; Schoffelen *et al*., 2017).

Very similar paradigmatic choices have been made in electroencephalography (EEG) and MEG studies that have examined changes in local cortical activation: fully naturalistic paradigms have been used to investigate speech perception (Gross, Hoogenboom, *et al*., 2013; Alexandrou *et al*., 2017, 2018; Koskinen *et al*., 2020) whereas continuous reading with eye movements has been investigated almost exclusively at the sentence level (Vignali *et al*., 2016; Metzner *et al*., 2017a; Loberg *et al*., 2019; Pfeiffer *et al*., 2020). Largely, these differences stem from the differences in artefactual signals between speech perception and reading. Eye movements that are an integral part of natural reading (Rayner, 1998; Metzner *et al*., 2017b) induce severe artefacts to electrophysiological signals recorded with MEG and EEG (Dimigen *et al*., 2011). While such artefacts can be detrimental for accurate estimation of local cortical activity, they are even more problematic for studying cortico-cortical connectivity. We have recently tested and evaluated distinct approaches for removing ocular artefacts from continuous natural reading consisting of full-page texts (Mäkelä *et al*., 2022). Such reading induces several types of ocular artefacts, including blinks, forward and backward saccades and return-sweeps related to line changes. By utilizing either Adaptive Mixture ICA (AMICA) (Palmer *et al*., 2008) or an approach combining multiple blind source-separation techniques (Jutten & Herault, 1991; Belouchrani *et al*., 1997; Hyvärinen, 1999), also such richer artefactual patterns can be removed from the MEG data (Mäkelä *et al*., 2022), enabling the study of modulation of neural connectivity during natural reading. In this study, we applied AMICA to suppress the eyemovement artefacts as using a single method allows them to be identified in a more straightforward manner.

Overall, the existing literature on the neural underpinnings of reading builds on controlled event-related paradigms and involves measures of both cortical activity and connectivity. This literature presents a complex pattern of cortical structures and neural frequencies that support specifically reading and language processing as well as those that are necessary for processes such as working memory and attention that are critical for reading beyond the word level. As regards the possible cortical structures and their interconnectivity, regions particularly within the occipital, parietal, middle temporal and inferior frontal cortex have been shown to support reading or attentional control and working memory (Pugh *et al*., 1996; Fiez & Petersen, 1998; Jobard *et al*., 2003; Owen *et al*., 2005; Salmelin & Kujala, 2006; Jensen *et al*., 2007; Petersen & Posner, 2012; Nacher *et al*., 2013; Schoffelen *et al*., 2017). In the spectral domain, modulation of cortico-cortical coherence and directed neural interactions have been observed in speech perception and production as well reading studies across a wide range of frequencies including the theta, alpha, lowand high-beta as well lowand high-gamma bands (Kujala *et al*., 2012; Liljeström, Kujala, *et al*., 2015; Saarinen *et al*., 2015; Schoffelen *et al*., 2017; Liljeström *et al*., 2018). Notably, these frequency bands largely overlap with bands that have been shown to be central in supporting feedforward and feedback influences in visuospatial attention (Bastos *et al*., 2015; Michalareas *et al*., 2016). Thus, previous research has reported a rich set of cortical structures and distinct spectral connectivity patterns that are required in reading connected text and in visuospatial attention. However, it has not been addressed what kind of connectivity between brain regions would most critically support natural reading that incorporates, e.g., moment-to-moment linguistic processing and mutual influence between neural systems that support general cognitive and more specific language-related processing.

Here, our goal was to study the cortex-wide integration of information during naturalistic reading by quantifying the modulation of long-range cortico-cortical connectivity patterns during reading versus visual scanning of full-page texts. We expected that naturalistic reading would be associated with increased coherent interactions compared to the scanning task, reflecting integration of information across different neural systems, particularly those supporting lexical access and sentence-level unification inherent to continuous reading.

## 2 Methods

### 2.1. Participants and experimental Design

Eighteen right-handed Finnish-speaking subjects with normal or corrected-to-normal vision participated in the study. Five of the subjects were left out of the analyzed data as these subjects failed to comply to the task instructions in the visual scanning task (substantially different eyemovement patterns between the scanning and reading tasks or text being read during the scanning task as indicated by the post-experiment questionnaire), leaving 13 subjects (7 men, 6 women; age 20-50 years, mean 25.4; SD 8.3 years) in the final cohort. None of the subjects reported a history of neurological abnormalities or psychiatric disorders. Informed consent was obtained from all subjects, in agreement with a prior approval of the local Ethics Committee (Hospital district of Helsinki and Uusimaa). The study was conducted in accordance with the guidelines of the Finnish National Board on Research Integrity.

### 2.2. Experimental Design

The subjects were presented with a naturalistic reading task, as well as a visual scanning task to serve as a baseline condition for comparison. In the reading task, the subjects were instructed to read the texts in their usual way. In the scanning task, they were told to search for horizontally flipped letters “a” and “e”, mimicking eye-movements in normal reading. The instruction was to read or scan each text only once from left to right. This type of a scanning task induces eye movement patterns that are very similar to normal reading (Vitu *et al*., 1995; Rayner & Fischer, 1996). The subjects were divided into two groups. The first group read the texts that the second group scanned, and the second group read the texts that the first group scanned. In both tasks, the stimuli were eight different 3-page long texts picked from various Finnish-language novels and essays, slightly modified to be suitable for our study. The subjects changed the page of the text by lifting their right index finger. Each page of text comprised eight lines. There were on average 87.8 words on each page (SD 6.4), and 11.0 words on each line (SD 1.5). In the scanned texts, the locations (line and position in a word) of the flipped letters were randomized for each page of text. There were three flipped letters in 25%, four in 38%, five in 23% and six in 15% of the scanned pages. Out of the flipped letters, 80 % were flipped e’s and 20% flipped a’s. Before each text, a page with a yellow text “read” or a blue text “scan” was presented to inform the subject about the upcoming task. Each text was followed by 1-2 questions (12 in total for both reading and visual scanning) to measure participants’ comprehension of the text (reading task) or to address their search for flipped letters (scanning task: “Were there more than one type of flipped letters in the text?”). The number of scanned texts with a single type (52%) or two types (48%) of flipped letters was balanced, and for the scanned texts with two types of flipped letters the second type was presented at the end of the page to motivate the subject to scan the whole text. The subjects responded verbally, and their answers were written down for further analysis. The whole experiment was followed by a surprise questionnaire to test how the subjects had processed the semantic content in the two tasks. This surprise quiz consisted of 40 sentences in a randomized order, each beginning with the phrase “During the experiment I saw a text in which …”. The subjects were asked to answer “Yes”, “No” or “I don’t know”. 10 sentences described reading tasks, 10 scanning tasks and 20 sentences were non-related.

### 2.3. MEG recordings and preprocessing

The subjects’ brain activity was recorded with a 306-channel MEG system (Elekta-Neuromag VectorView, Helsinki, Finland), band-pass filtered at 0.03-200 Hz and sampled at 600 Hz. Vertical and horizontal electro-oculogram (EOG) signals were recorded for monitoring blinks and saccades. Anatomical MRIs were obtained with a 3T General Electric Signa system (Milwaukee, USA). Eye movements during the experiment were monitored with an Eye Link 1000 eye tracker (SR Research Ltd.; Mississauga, ON, Canada) using a sampling rate of 1000 Hz. The MEG data were preprocessed using the temporal extension of the Signal Space Separation (SSS) method (Taulu & Simola, 2006). Subsequently, the eye-movement artefacts (blinks and saccades) were removed using AMICA (Palmer *et al*., 2008) and in-house code for selecting the artefactual components based on examination of the component time series and evaluation of the sensor topographies of the components (Mäkelä *et al*., 2022). In the subsequent analyses, data from only the 204 gradiometers were used as they are less prone to pick up signals from distant sources and external artefacts than magnetometers. The effectiveness of the artefact removal pipeline was evaluated by examining the mean power spectra across the two tasks for the MEG data with and without AMICA preprocessing. This was conducted both for 12 different sensor groups distributed across the cortex (12-20 sensors per group) and for MEG data averaged across all sensors.

### 2.4 Analysis of eye-movement patterns

To identify the possible differences in eye movements between the reading and scanning conditions we quantified and examined the properties of the fixations and saccades occurring during the two tasks. Specifically, we determined, for each subject, the mean duration of the fixations and saccades, the length of the forward saccades and the number of backward saccades for both tasks. The possible differences between the tasks were evaluated with paired-samples t-tests (p < 0.05).

### 2.5. MEG data analysis

Long-range cortico-cortical coherence was determined using Dynamic Imaging of Coherent Sources (DICS, Gross *et al*., 2001; Kujala *et al*., 2007) in six different frequency bands (4–7, 8–13, 13–20, 20–30, 35–45, 60–90 Hz). In the coherence estimation, one subject’s brain was first fitted with a surface-based grid (9-mm spacing along the surface of the cortex) in MNE (Gramfort *et al*., 2014) that was transformed into the other subjects’ anatomies, leading to spatially equivalent sampling across the subjects. Grid points that were further than 7 cm from the closest MEG sensor were excluded from the analysis. Coherence estimation was based on an approach that utilizes the numerical maximization of coherence across a set of source orientation combinations (Liljeström, Kujala, *et al*., 2015; Liljeström, Stevenson, *et al*., 2015; Liljeström *et al*., 2018). Specifically, the source current orientations for each connection were determined by identifying the orientation combination that maximized the mutual coherence between the two sources. This was done separately for the two experimental conditions by using 50 regularly spaced tangential orientations for both sources. Coherence was estimated across all grid point combinations for connections that were longer than 4 cm; the length threshold was set to avoid spurious coherence detection due to field spread effects that influence especially short-range connections (Schoffelen & Gross, 2009; Liljeström, Kujala, *et al*., 2015). These connections were then averaged across cortical parcel pairs. The set of parcels of interest was based on the automatically labeled anatomical parcellation consisting of 35 regions per hemisphere (Desikan *et al*., 2006). Sixteen of these regions that have been demonstrated to be involved in reading or other cognitive processes (Pugh *et al*., 1996; Fiez & Petersen, 1998; Jobard *et al*., 2003; Owen *et al*., 2005; Salmelin & Kujala, 2006; Jensen *et al*., 2007; Petersen & Posner, 2012; Nacher *et al*., 2013; Schoffelen *et al*., 2017) were selected for the analysis (see Figure 1). The parcel-level all-to-all coherence estimation was done within each hemisphere, and the modulation of cortico-cortical coherence between reading and scanning of text was evaluated using Wilcoxon signed rank test, corrected for multiple comparisons using false discovery rate (FDR) correction. For connections showing significant modulation of cortico-cortical coherence between the tasks, we examined whether the magnitude of the coherence modulation would be correlated with the results of the surprise questionnaire (accuracy difference between the tasks), using Spearman’s rho (p < 0.05).

**Figure 1.**
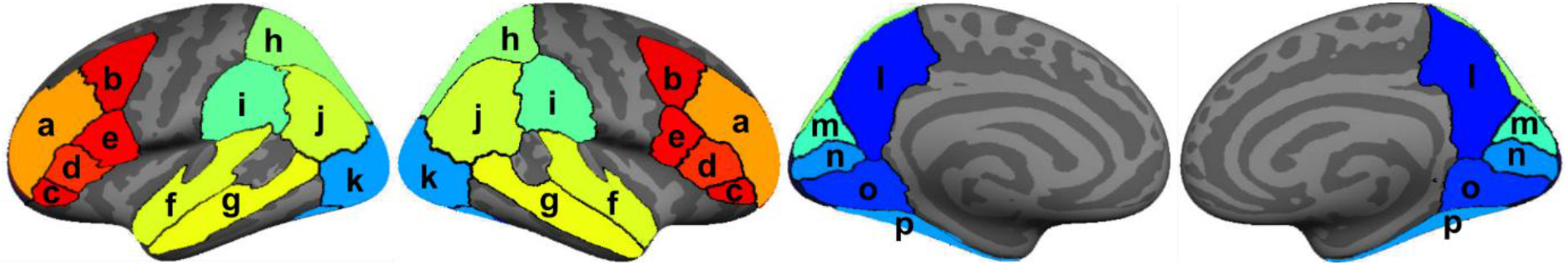
Parcels selected for the analyses. Bilateral rostral middle frontal cortex (a), caudal middle frontal cortex (b), pars orbitalis (c), pars triangularis (d), pars opercularis (e), superior temporal gyrus (f), middle temporal gyrus (g), superior parietal gyrus (h), supramarginal gyrus (i), inferior parietal gyrus (j), lateral occipital cortex (k), precuneus (l), cuneus (m), pericalcarine cortex (n), lingual gyrus (o) and fusiform gyrus (p). Identical sets of parcels were used in the left and right hemisphere.

For connections showing significant increase of cortico-cortical coherence between the reading and visual scanning tasks, we also estimated whether the modulation of inter-areal synchrony would be associated with increases in directed neural interactions between the two conditions. Here, we used the multivariate Granger causality toolbox (MVGC, Barnett & Seth, 2014) to estimate the frequency-domain Granger Causality (GC) within the same frequency bands that were used in the analysis of cortico-cortical coherence (4–7, 8–13, 13–20, 20–30, 35–45, 60– 90 Hz). Common beamformer weights across conditions were first obtained with DICS in the 1–45 Hz band. Here, the averaged cross-spectral density matrix across the reading and scanning tasks was used to estimate the source orientation that maximized the signal power at each grid point. This orientation was used to estimate the beamformer weights at each grid point which were then averaged within each parcel of interest to obtain a single timeseries per subject and condition that represented the neural activity at each parcel. These timeseries enabled the application of uniform autoregressive modelling and GC analysis across all frequency bands of interest. Here, for each tested connection between parcel pairs, the timeseries were segmented into 2-s long trials. Linear trends were removed from these trial-level timeseries, and the timeseries were also demeaned and subjected to first-order differencing. Akaike and Bayesian information criteria were used to determine the optimal model order for GC estimation. To facilitate comparison of the GC values across conditions, we used for all estimates the same model order (30) that fell in the middle of the estimated model orders across subjects, the two information criteria and all connections. As the computed GC values represent biased estimates of the true GC of the underlying neural processes, we applied an unbiasing approach that allows for contrasting the GC values across the experimental conditions (Barrett *et al*., 2012). Here, for each connection and both conditions, the extracted 2-s long trials were randomly paired 200 times across the two parcels forming the connections. As each of these randomized pairs has a true GC value of zero, the process yields an approximation of a GC null distribution. By subtracting the mean values across the 200 obtained values from the original GC estimate, we then acquired unbiased GC values that can be subjected to statistical contrasting between the reading and scanning tasks. In the contrasting, we examined whether increases in cortico-cortical coherence for reading vs. scanning would be associated with increased directed interactions between the reading and visual scanning task separately for both directions of each connection (i.e., from parcel 1 to 2 and from parcel 2 to 1) using one-tailed Wilcoxon signed rank tests.

## 3 Results

The questions following each text were answered with 87±13% (mean±SD) accuracy in the reading and 80±8% accuracy in the scanning task. There was no difference between the reading and visual scanning task in the task performance, as determined using Wilcoxon signed rank test (p = 0.195). The surprise questionnaire presented after the experiment, in turn, indicated that the subjects had processed the semantic content during the reading task (accuracy 87±8%) better (Wilcoxon signed rank test, p = 0.00024) than during the scanning task (accuracy 27±20%).

The reading and scanning conditions were associated with closely matched eye-movement patterns for all fixation and saccade measures: fixation duration 237±34 ms (mean±SD) for reading and 231±31 ms for scanning; saccade duration 30±5 ms for reading and 29±3 ms for scanning; length of forward saccades 60±13 units for reading and 69±17 for scanning; number of backward saccades 846±335 for reading and 781±402 for scanning. The statistical analysis revealed a significant difference in the length of the forward saccades between the tasks (p = 0.0027) whereas no significant effects were detected for the other three measures (p > 0.1).

As the eye-movement patterns and the artefactual magnetic field patterns associated with them were not identical during the reading and scanning tasks, we applied a procedure based on the use of AMICA to suppress the eye-movement artefacts. Figure 2 shows the influence of the artefact suppression for MEG data averaged across the two conditions. The areal average spectra (Figure 2A) show that artefacts were suppressed primarily in the frontal and anterior temporal MEG sensors. The quantification of these effects across all MEG sensors (Figure 2B) revealed that the suppression was in the range of 1-10% across different frequencies, with a marked emphasis on the frequencies below 10 Hz, a ca. 1% effect across most of the spectrum, and a ca. 2% effect for specific, narrow frequency bands (11-13 Hz and 22-24 Hz).

**Figure 2.**
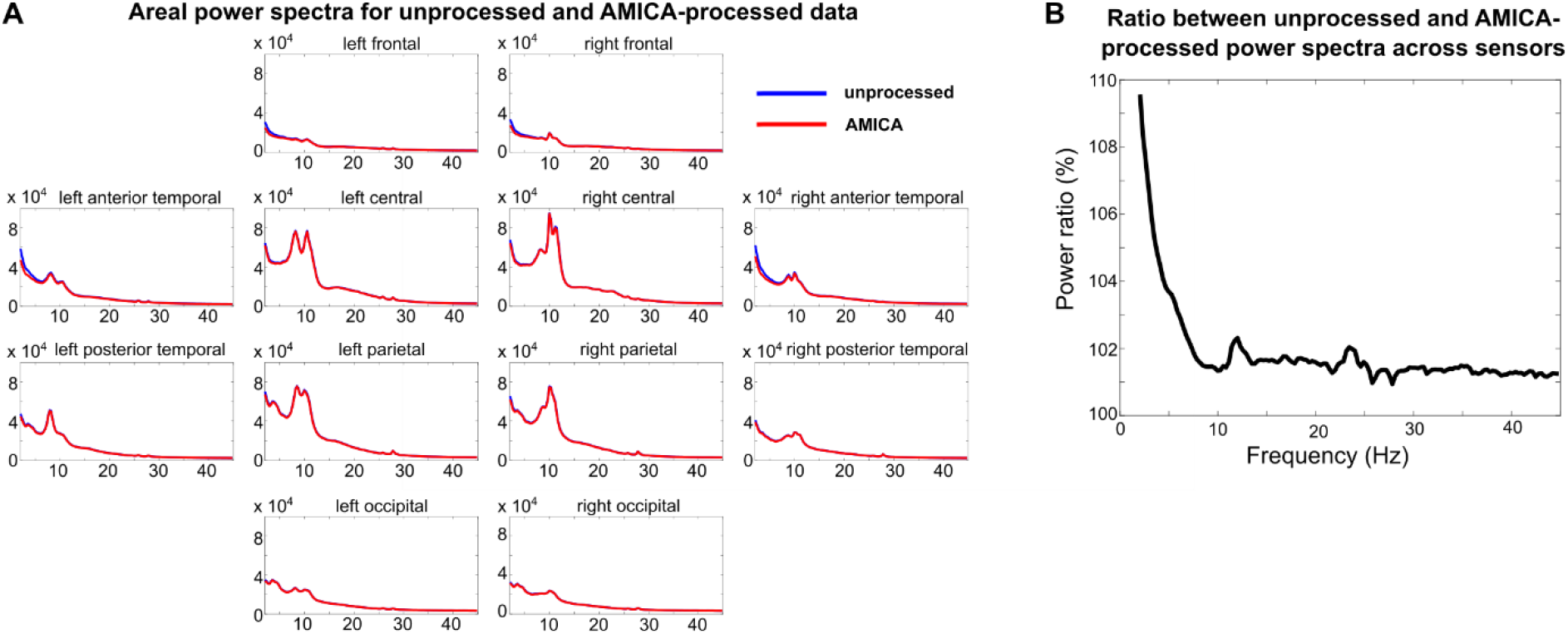
Comparison between sensor-level power spectra averaged across subjects and the two conditions for data with and without AMICA-based preprocessing. A) Areal averages in different MEG sensor groups. The x-axis portrays the frequency (Hz) and the y-axis the magnitude (fT^2^/cm^2^) of the spectra. B) Ratio between the power spectra averaged across all MEG sensors for the unprocessed and AMICA-processed data.

**Figure 3.**
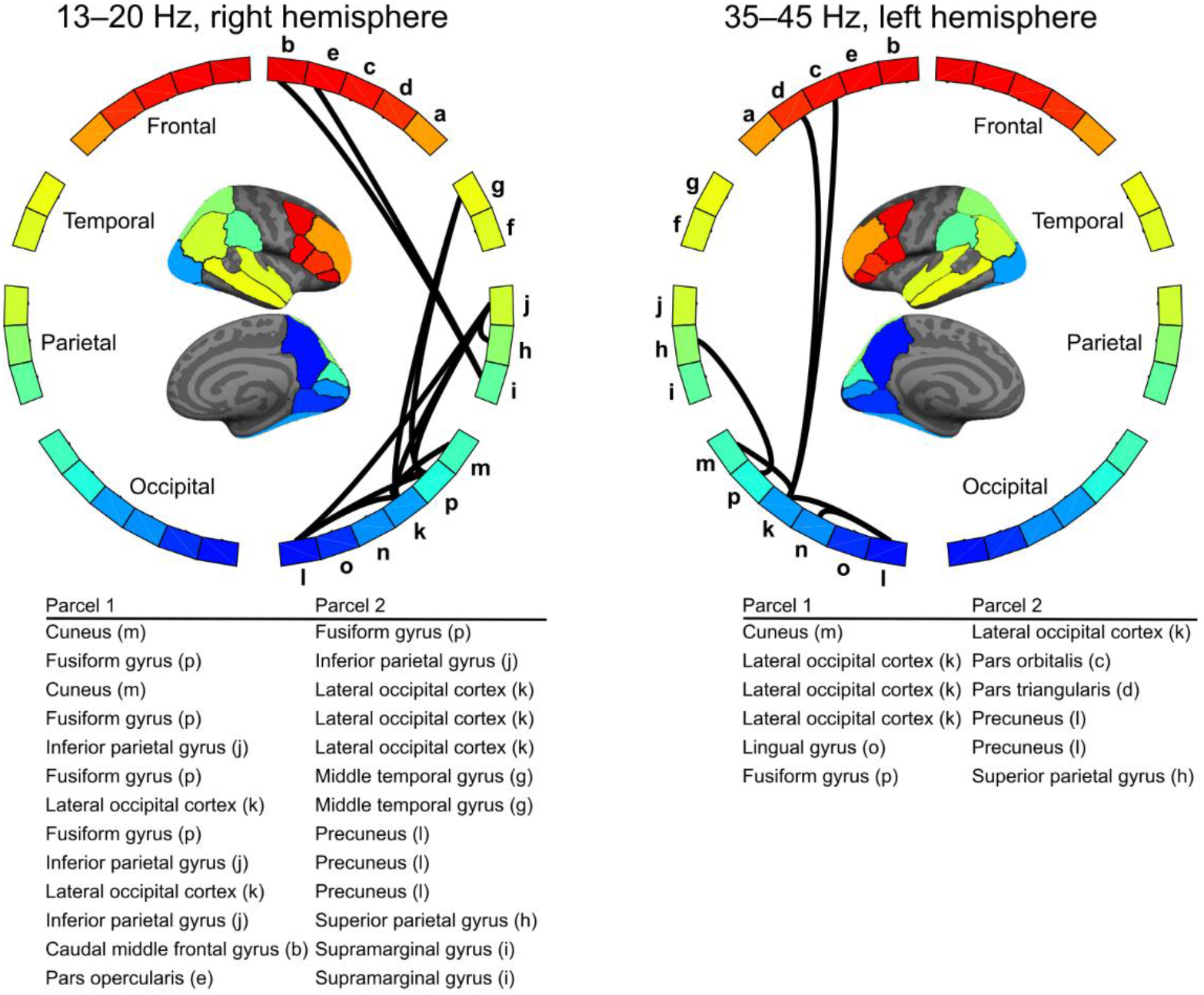
Parcel-level connections showing higher cortico-cortical coherence for reading vs. visual scanning of text (p < 0.05, FDR). Statistical testing was conducted separately within each hemisphere in six different frequency bands (4–7, 8–13, 13–20, 20–30, 35–45, 60–90 Hz). Significant modulation of cortico-cortical coherence was detected only in the 13–20 Hz and 35–45 Hz bands. See Fig. 1 for all parcel labels.

The estimation of all-to-all coherence in the six frequency bands of interest across the 16 parcels per hemisphere revealed significant (p < 0.05, FDR, Wilcoxon signed rank test)

modulation of cortico-cortical coherence in the low beta-band (13–20 Hz) and the low gamma-band (35–45 Hz) between the reading and scanning tasks (see Figure 2). For all detected connections, coherence was higher during reading than scanning of text. Significant effects were detected exclusively in the right hemisphere for the 13–20 Hz band and in the left hemisphere for the 35–45 Hz band. In the right hemisphere, modulation of coherence was detected within the occipital and parietal cortices as well as between the occipital and parietal, occipital and temporal, and parietal and frontal cortex. In the left hemisphere, coherence was modulated within the occipital cortex and between the occipital cortex and the parietal as well as the frontal cortex. The testing for possible correlation between the amount of coherence increase and accuracy difference in the post-experiment questionnaire, across the tasks, did not yield and significant results (p > 0.05 for all 19 connections).

The evaluation of Granger Causality across the connections showing significant modulation of coherence revealed that directed influences as quantified using the unbiased GC estimates were increased for reading vs. visual scanning across all the examined frequencies except the 60–90 Hz band (p < 0.05, one-tailed Wilcoxon signed rank test). The reading vs. the scanning task was associated with higher GC from the left Lateral occipital cortex to the left Precuneus at 20–30 Hz and to the left Pars triangularis at 4–7 Hz (Figure 4). Within the right hemisphere, higher GC for reading vs. scanning was detected from the Inferior parietal gyrus to the Superior parietal gyrus at 8–13 Hz and to the Lateral occipital cortex at 13–20 Hz, from the Fusiform gyrus to the Middle temporal gyrus at 8–13 Hz, 13–20 Hz and 20–30 Hz, from the Middle temporal gyrus to the Fusiform gyrus at 35–45 Hz, and from the Fusiform gyrus and Precuneus to the Lateral occipital cortex at 35-45 Hz.

**Figure 4.**
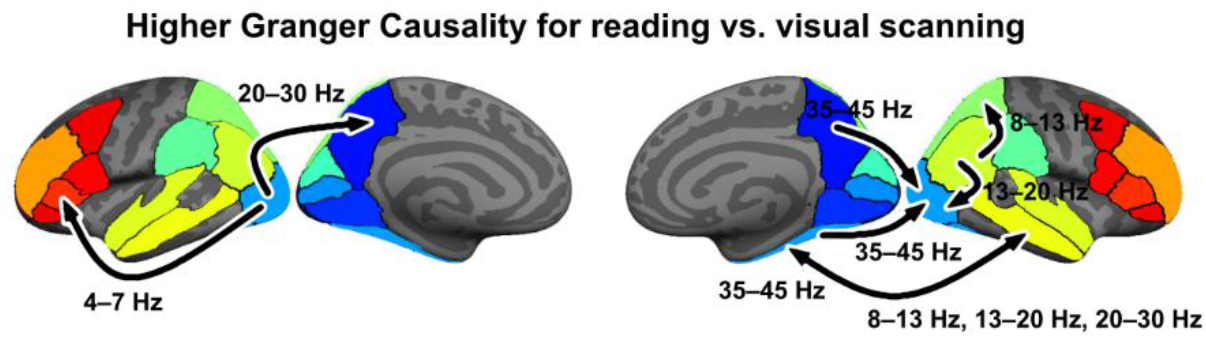
Connections showing significantly higher Granger Causality for reading than scanning of text. The arrows indicate the direction of the Granger Causality effects with the arrowhead marking the receiving end of the influence. The frequency-band where the effects were detected are marked next to the connections. Note that for the GC effects between the right Fusiform gyrus and Middle temporal gyrus bi-directional effects were detected, with the Middle temporal cortex influencing the Fusiform gyrus at 35–45 Hz and the Fusiform gyrus influencing the Middle temporal gyrus in several neighboring frequency bands (8–13 Hz, 13– 20 Hz and 20–30 Hz).

## 4 Discussion

Here, we demonstrate that by combining advanced artefact rejection protocols and closely matched experimental conditions (Vitu *et al*., 1995; Rayner & Fischer, 1996), the modulation of cortico-cortical connectivity related to naturalistic reading can be investigated with electrophysiological neuroimaging. Specifically, we show that both coherent coupling and directed influences between brain regions are modulated between reading and visual scanning of full-page texts in the beta- and gamma-bands for phase synchronization as well across a broad range of frequencies for directed neural interactions. The observed modulations show that reading is associated with increased cortico-cortical coherence between brain regions compared to text scanning in the beta and gamma bands. This increased connectivity presumably supports the large-scale integration of information across cortical networks that is crucial for linguistic processing of written text. This notion is supported by the finding that the subjects could significantly more accurately answer the surprise questionnaire following the experiment when the questions pertained to the texts that were read instead of scanned. Possible differences in the content of the read and scanned texts, as such, is not a likely explanation for the modulation of coherence as the texts were counter-balanced across subjects. Among the connections that showed increased inter-areal coherence associated with reading, we additionally observed, especially in the right hemisphere, increased directed neural interactions for reading vs. scanning across a broad range of frequencies. Together, our findings demonstrate the role of long-range neural interactions in naturalistic reading and highlight the role of synchronized beta- and gamma-band oscillations and spectrally distributed directed neural interactions in supporting the reading process.

### 4.1. Topography of modulations of cortico-cortical connectivity in natural reading

Modulation of coherent coupling between the reading and visual scanning tasks was observed prominently within the occipital cortex, likely reflecting the task-induced requirement of forming words from the visual input during reading vs. detection of inverted letters during scanning. Such patterns were observed both within the left and the right hemisphere. These neural interactions align with previous findings that increased memory load is associated with increased inter-areal synchrony within the occipital cortex (Palva, Monto, Kulashekhar, *et al*., 2010) and that directed interactions from the dorsal to the ventral visual stream support task performance during working memory (Popov *et al*., 2018). Increased cortico-cortical coherence was also detected within the parietal cortex, between the right Superior and Inferior parietal parcels, and cross-lobar connectivity modulation in both hemispheres between the occipital and parietal cortices. These modulations likely reflect the differences between the reading and visual scanning tasks in directing visuospatial attention and in integrating top-down influences with bottom-up information across the visual hierarchy (Buffalo *et al*., 2010; Doesburg *et al*., 2016; Michalareas *et al*., 2016).

In the left hemisphere, cortico-cortical coherence between the occipital and inferior frontal cortex was significantly higher during the reading than the scanning task, whereas within the right hemisphere we observed a corresponding increase between the parietal and frontal cortex. These findings suggest that that the processing of written text in the left hemisphere is supported by the more direct unification of words within the language network (Kujala *et al*., 2007; Wang *et al*., 2018), whereas in the right hemisphere the modulation of neural interactions reflect a multi-stage hierarchy in integrating visual information into linguistic content.

Similar observations have been made in previous studies that have shown increased coherence for isolated words compared to symbols in the left hemisphere between the occipital and frontal cortices (Liljeström *et al*., 2018) and increased inter-areal coupling between the right parietal and frontal cortex associated with visual comparison (Saarinen *et al*., 2015). A notable difference compared to previous studies on neural interactions during reading (Kujala *et al*., 2007, 2012; Schoffelen *et al*., 2017; Liljeström *et al*., 2018) is the relatively small amount of observed connectivity from the temporal cortices to other brain regions in the present study. One likely reason for this is that whereas previous studies have focused on demonstrating the existence of neural interactions during reading (Schoffelen *et al*., 2017) or used stimuli with varying linguistic content (Liljeström *et al*., 2018), we were specifically interested in connections that were modulated across experimental conditions that had the same text stimuli but during which the tasks of the participants were different. Our findings suggest that some of the previously reported connectivity from the temporal cortices to other regions are primarily dependent on the content of the language stimuli rather than the unification of the words into a coherent whole, as such (Kujala *et al*., 2007; Schoffelen *et al*., 2017; Liljeström *et al*., 2018). However, in both hemispheres, we observed increased coupling between the temporal and occipital cortex for reading compared to scanning, suggesting that the connectivity of the temporal cortices is important for semantic processing of words.

We detected modulation of cortico-cortical connectivity in just two of the six frequency bands that we examined in the study. Moreover, within each hemisphere, only individual bands showed significant results. The observed network patterns are thus likely to represent only a portion of the full range of cortico-cortical connectivity necessary for supporting natural reading. Accordingly, the present data do not allow us to make strong inferences on the distinct roles of the two hemispheres in natural reading or the possible differences in shorter- and longer-range neural connectivity across the hemispheres. One interesting notion, however, was obvious in the results from the Granger Causality analysis. Within the left hemisphere, we detected only bottom-up directed neural interactions whereas in the right hemisphere also top-down influences were seen. It is thus possible that the connectivity patterns within each hemisphere play distinct roles in supporting more automatic hierarchical processing (left hemisphere) vs. updating the linguistic expectation set by the preceding word sequences (right hemisphere). However, as the set of connections showing modulation of directed neural interaction was very sparse, this interpretation necessitates further studies that allow the examination and identification of such modulations across a much broader range of connections.

### 4.2. Role of inter-areal synchrony and directed neural interactions in reading and scanning of text across different frequency bands

We detected modulation of cortico-cortical coherence in the beta and gamma bands that have been previously shown to be vital in supporting network-level neural dynamics in visual attention as well as picture naming and reading (Bosman *et al*., 2012; Kujala *et al*., 2012; Bastos *et al*., 2015; Liljeström, Kujala, *et al*., 2015; Schoffelen *et al*., 2017; Wang *et al*., 2018). Specifically, these frequency bands have been shown to support both feedforward and feedback influences in visuospatial attention, phonological processing, and sentence unification as well as to be involved in anticipatory unification and in facilitatory and suppressive roles during speech production. It has also been proposed that the beta and gamma bands have distinct functions in word prediction from cumulative semantic interpretation and in accommodating words to the cumulative interpretation (Meyer, 2018). Notably, the detected modulations showed exclusively effects where the inter-areal coupling was stronger for reading than scanning. At a first glance, these findings align with the existing evidence that has shown the importance of these oscillations in both bottom-up and top-down neural interactions (Bosman *et al*., 2012).

However, when we examined the directed neural interactions with Granger Causality, the alignment between our and previous findings was more ambiguous. First, we did not detect effects of bottom-up directed influences in the gamma-band that have been suggested to be central especially in visual attention (Bosman *et al*., 2012; Bastos *et al*., 2015; Michalareas *et al*., 2016). Instead, we observed top-down influences within the gamma-band and both bottom-up and top-down influences in the beta-band. In the lower theta and alpha bands, we detected exclusively bottom-up directed neural interactions. These findings agree conceptually with previous MEG work on reading where wide-spread directed neural interactions were detected across several other frequencies but not the gamma band, an effect argued to stem from the weakness of the gamma-band responses in language compared to visual processing (Schoffelen *et al*., 2017). Thus, it is possible that the systematic detection of modulation of directed, bottom-up neural interactions within the gamma-band in reading may necessitate a fairly high signal-to-noise ratio that is available with intracranial recordings but typically not afforded by MEG (Perrone-Bertolotti *et al*., 2012). Accordingly, the connections within the gamma band that can be identified with MEG during reading may represent isolated phenomena and not be informative on whether the gamma-band interactions would, in fact, be more involved in bottom-up than top-down processes.

The second slight misalignment between our and previous studies utilizing Granger Causality is the role of top-down beta-band neural interactions. Previous studies on the visual system have systematically indicated that beta-band directed interactions are central in conveying top-down influences (Bastos *et al*., 2015; Michalareas *et al*., 2016), a finding that has been largely replicated for sentence-level reading (Schoffelen *et al*., 2017). Here we detected both bottom-up and top-down directed interactions within the beta band, without a clear directional dominance. The only spectrally systematic observation on directionality in the present study was that directed interactions in the lower frequency bands (theta, alpha) band were exclusively bottom-up in nature, an observation that aligns with previous literature (Bastos *et al*., 2015; Schoffelen *et al*., 2017). In general, our findings suggest that naturalistic reading may involve spectrally more wide-spread inter-areal coupling than shown previously (Kujala *et al*., 2007), possibly reflecting efficient integration of linguistic information across large-scale networks.

### 4.3. Methodological considerations

In the present study we contrasted natural reading of full-page texts to visual scanning of identical texts for flipped letters. This choice imposes several potential confounding factors and limitations to the observed results and their interpretation. First, while the eye-movement patterns during reading and scanning were closely matched, the subjects made on average longer forward saccades during scanning than reading. As the eye-movement patterns were not identical between the conditions, it is possible that the observed findings stem partially from the differences in artefacts related to them. While we applied an AMICA-based procedure to suppress these artefacts and thus to minimize such effects, the possibility remains that the residual artefacts may induce observable differences in cortico-cortical connectivity across the

tasks. It should, however, be noted that the suppression of eye-movement artefacts affected the MEG spectra primarily in frequencies below 10 Hz as well as in individual narrower bands (11–13 Hz, 22–24 Hz). The frequency bands (13–20 Hz, 35–45 Hz) that showed modulation of cortico-cortical coherence were only marginally influenced by the artefact removal, also proposing that the effects observed in these bands are unlikely to be due to differences in the eye-movement artefacts. As for the Granger Causality findings, we applied an unbiasing procedure that accounts for systematic differences in the signal-to-noise ratio of the data. The findings should, thus, represent specifically modulation of directed interactions between signals instead of artefactual effects.

The contrasting of reading and scanning of identical texts also limits how detailed and specific questions could be addressed within the present study. Based on the collected behavioural data, we could determine that the subjects remembered the content of the texts better when they had read than scanned them. We cannot, however, determine whether this was due to differences in, e.g., lexical access of the words, unification at the sentence level or attention to words vs. individual letters. Accordingly, based on the present study it is not possible to infer the relationship between the observed connectivity modulation and specific linguistic subprocesses. Similarly, we cannot properly address how our findings align with predictions from different models of reading as they could be explained, e.g., by different emphasis on the lexical vs. non-lexical route (Coltheart *et al*., 2001) or on memory, unification or control in language processing (Hagoort, 2005). Future studies should parametrically manipulate these dimensions by contrasting reading of texts where only specific aspects differ. Such studies would also allow one to determine whether the modulations of cortico-cortical connectivity during continuous reading would reveal effects beyond those that can be observed in more traditional event-related paradigms, potentially linked to the integrative nature of moment-to-moment linguistic processing and eye-movement control that are inherent to natural reading (Rayner & Fischer, 1996).

In addition to the experimental design within the present study, also the relatively small sample size imposed limitations to the questions that could addressed and the interpretations that could be made based on the findings. As it was not possible to make specific hypotheses on which particular neural connections would be critical in natural reading, we analysed the modulations of cortico-cortical connectivity in an all-to-all manner. As a substantial proportion of the subjects failed to comply with the task instruction, the statistical power afforded by the subjects included in the study allowed us to detect only somewhat isolated neural effects. Specifically,

we observed significant modulations of cortico-cortical coherence in two frequency bands, one band in the left hemisphere and a different one in the right hemisphere. Thus, it is not possible to make in-depth interpretations on whether the network topography or connectivity across distinct oscillators would be fundamentally different across the hemispheres. Similarly, the limited statistical power within the study did not allow us to extend the analysis to the study of the role of interhemispheric connectivity. These questions would necessitate studies with larger sample sizes.

It should also be noted that within the present study we used all the collected MEG data, thus individually varying amounts of data in the two conditions, in estimation of cortico-cortical connectivity to minimize the variance and uncertainty of the estimates (Halliday *et al*., 1995). As we used a condition-specific beamforming approach that allows determining cortico-cortical coherence also for sources with weak activity levels (Liljeström *et al*., 2018), the potential signal-to-noise difference between the two conditions should not bias the magnitude of the coherence estimates. However, when investigating the amount of oscillatory power or when using different measures and approaches for determining cortico-cortical connectivity, such differences can bias the estimates across the conditions (Gross, Baillet, *et al*., 2013).

## 5 Conclusions

We demonstrate that coherent cortico-cortical connectivity and directed neural interactions across several frequencies are increased during naturalistic reading compared to visual scanning of full-page texts. In particular, our results show that reading is associated with increased coherent coupling across large-scale cortico-cortical networks in the beta and gamma band, and that natural reading depends also on spectrally rich, directed neural interactions. Our findings demonstrate how both synchronized and directed long-range neural connectivity can flexibly build on different frequency bands to support the large-scale integration of information necessitated by natural reading.

## Acknowledgements

This work was financially supported by the Academy of Finland (grants #255349 and #315553 to RS), the Sigrid Jusélius Foundation (grant to RS), the Finnish Cultural Foundation (grant to SM) and Aalto Brain Center. We thank Antti Jalava for contributing to the design and implementation of the experimental paradigm as well data collection.

